# Strain level variation in *Proteus mirabilis* chondroitin sulfate degradation kinetics and regulation by urea

**DOI:** 10.64898/2026.03.23.713754

**Authors:** Brianna M. Shipman, Serena Zhou, Benjamin Hunt, Vitus Brix, Ibrahim Salaudeen, Anna N. Evers, Brian S. Learman, Nicholas A. Dillon, Philippe E. Zimmern, Chelsie E. Armbruster, Nicole J. De Nisco

## Abstract

To establish infection, uropathogens must overcome several host defenses including the glycosaminoglycan (GAG) layer coating the apical surface of the bladder urothelium. GAGs are thought to protect against urinary tract infection (UTI) by serving as scaffolding sites for commensals, providing barrier function and preventing uropathogen adherence. However, the ability of uropathogens to degrade and utilize GAGs and the contribution of these activities toward UTI progression is largely unknown. We previously discovered that the uropathogen *Proteus mirabilis,* a common cause of catheter-associated UTI (CAUTI), degrades the GAG chondroitin sulfate (CS). In this study we sought to define the kinetics and regulation of CS degradation by diverse *P. mirabilis* strains clinically isolated from both recurrent UTI and CAUTI patients. We found variation in CS degradation kinetics between *P. mirabilis* strains and media types. However, CS degradation depended on conserved putative chondroitin sulfate ABC endo- and exolyases in all strains. Furthermore, we found that CS degradation in Pm123 was repressed by urea and that this repression was dependent on *P. mirabilis* urease activity. Complementation of the Pm123 endolyase into urea-insensitive HI4320 resulted in a urea-sensitive CS degradation phenotype suggesting functional differences between the Pm123 and HI4320 endolyases. Sequence alignment and structural modeling analysis identified two unique point mutations within the Pm123 endolyase that may contribute to urea sensitivity. Finally, unlike urea-insensitive *P. mirabilis* strains, Pm123 demonstrated attenuated swarming and loss of chondroitin endolyase activity had no effect on Pm123 virulence in a mouse CAUTI model. Our results suggest that the kinetics and regulation of CS degradation differ between *P. mirabilis* strains and in urea-sensitive strains, thus reduces the contribution of CS degradation to urovirulence during murine CAUTI.

**Importance:** This work demonstrates that the ability to degrade a common component of bladder mucosal surfaces, chondroitin sulfate, is a phenotype that is shared by multiple strains of the common catheter-associated UTI (CAUTI) pathogen *P. mirabilis*. We find that this activity is dependent on encoded chondroitin ABC endo- and exolyases, first described in *Proteus vulgaris*. Additionally, we discovered that for *P. mirabilis* strain Pm123, degradation of CS is negatively regulated by the presence of urea, a major component of urine. The repression of CS degradation by urea is dependent on the activity of the *P. mirabilis* urease enzyme, which breaks down urea producing ammonia which raises pH. We found expression of the Pm123 CS endolyase was sufficient to confer a urea-sensitive CS-degradation phenotype and identified two unique mutations within the Pm123 enzyme that may contribute to urea sensitivity. Finally, we find that while CS-degradation plays a role in progression and severity of murine CAUTI model in urea-insensitive *P. mirabilis,* there was not significant difference in CAUTI outcomes between the urea-sensitive Pm123 wild-type and chondroitinase knockout strains. This study represents a major step forward in understanding the diversity of CS degradation activity and regulation among clinical strains of the critically important CAUTI pathogen *P. mirabilis* as well as its contribution to urovirulence.

## Introduction

Urinary tract infection (UTI) is generally categorized as either complicated or uncomplicated with complicated UTI being characterized by the presence of either physical abnormalities of the urinary tract or impairment of the host immune system (1, 2). One of the most common forms of complicated UTI is catheter associated UTI (CAUTI) representing approximately 80% of complicated UTI cases (3, 4). CAUTI is also heavily associated with hospital acquired infections (HAI) with estimates placing CAUTI as representing up to ∼40% of HAIs with incidence increasing as duration of catheterization progresses (4, 5). Also, while female sex is a major risk factor for uncomplicated UTI, CAUTI rates are similar in both sexes with some studies even finding a higher incidence in males (3, 5–8).

CAUTI also carries with it differing risks in regard to the causative urinary pathogens with bacteria less frequently found in uncomplicated UTI, such as *Proteus mirabilis* and *Enterococcus faecalis,* making up a much larger proportion of CAUTI cases (9–11). This difference in pathogen breakdown is due in part to the unique route of invasion in CAUTI, as the presence of the catheter surface serves as a substrate that allows for invasion of highly motile or adherent species; for example, studies have demonstrated efficient movement of *P. mirabilis* over the surface of multiple types of urinary catheters due to the swarming motility that is characteristic of the species (12, 13).

*P. mirabilis* is a Gram-negative opportunistic pathogen that is heavily associated with CAUTI, representing as much as ∼44% of cases (9, 10, 14). *P. mirabilis* is a particularly difficult uropathogen to treat as it is known to possess multiple clinically relevant virulence mechanisms including a urease enzyme that breaks down urea into ammonia for use as a nitrogen source (15–19). Additionally, *P. mirabilis* exhibits robust swarming motility whereby this species differentiates into highly motile swarmer cells that promote invasion of the lower and upper urinary tracts (13, 17, 20). *P. mirabilis* CAUTI also presents patients with a unique set of potential risk factors in terms of disease progression as its urease activity raises urinary pH causing struvite (MgNH_3_PO_4_) and apatite (CaPO_4_) crystalline biofilms to form, which result in encrustation and blockage of the catheter (21–27).

In addition to the aforementioned virulence mechanisms, prior work in our lab has revealed that *P. mirabilis* degrades the glycosaminoglycan (GAG) chondroitin sulfate (CS), the primary GAG in urine that comprises the GAG layer on the apical surface of the bladder epithelium (28–30). This GAG layer is thought to protect the bladder epithelium by acting as a physical barrier and preventing adherence of uropathogens thereby contributing to their expulsion from the urinary tract through the sheering force of urine (30–33). Additionally, studies have shown GAG degradation to be an important virulence mechanism in other species including *Streptococcus agalactiae* (Group B *Streptococcus*). *S. agalactiae* degrades the GAG hyaluronic acid (HA) thereby enabling immune evasion with the resulting HA disaccharides blocking TLR2/4 signaling as well as promoting ascending infection and pre-term birth (34, 35). There has also been recent evidence that GAG degradation may play a role in CAUTI as it was found that *E. faecalis* encodes two putative hyaluronidases termed HylA and HylB that when inactivated result in reduced bladder invasion and dissemination into the bloodstream (HylA) or a reduction in kidney invasion (HylB). These results suggest that GAG degradation may be involved in progression of ascending *E. faecalis* CAUTI and associated with worse disease outcome (36). However, only one of the two predicted hyaluronidases, HylB, has been shown to functionally degrade GAGs (HA and CS), and the exact mechanisms or signals underlying the *E. faecalis* hyaluronidase activity are unknown, indicating there is still much to be understood in how GAG degradation may impact CAUTI progression (36).

Our previous work identified CS degradation activity in three strains of *P. mirabilis* (28). Given these findings and what is known about GAG degradation in other pathogens, we hypothesized that CS degradation by *P. mirabilis* might be a clinically relevant virulence mechanism in the progression of CAUTI. To this end we characterized CS degradation kinetics and regulation in several *P. mirabilis* strains, including six isolated from the urine of postmenopausal women with either recent recurrent UTI history or active recurrent UTI, as well as the CAUTI type strain (HI4320) originally isolated from the urine of an elderly woman with a long-term indwelling urinary catheter. Herein, we report significant strain-level variation in the kinetics of CS degradation and utilization among *P. mirabilis* isolates and media types. We show that CS degradation activity is lost in genetically tractable clinical isolates following targeted deletion of genes encoding putatitive chondroitin ABC endolyase and exolyases proteins with near perfect homology to the well-characterized *Proteus vulgaris* enzymes (37–40). In addition to significant variation between CS degradation kinetics between strains, we identify a unique *P. mirabilis* strain, Pm123, for which GAG degradation is repressed by urea, a phenotype that is likely due to differences in protein structure or stability. Additionally, we report evidence that for urea insensitive strains, CS degradation may play a role in worsened disease outcomes in a mouse model of CAUTI.

## Results

### Isolation and phylogenetic analysis of *P. mirabilis* clinical urinary isolates

Six *P. mirabilis* strains were isolated from the urine of postmenopausal women visiting the urology clinic at UT Southwestern Medical Center. All strains were isolated from either women with recent recurrent UTI (rUTI) history but no active UTI (rUTI remission) or current active rUTI (rUTI relapse) (**Figure 1A**). In these urine samples, *P. mirabilis* was frequently cultured alongside multiple other pathogenic and commensal species including members of the genera *Actinobaculum*, *Actinomyces*, *Aerococcus*, *Alloscardovia*, *Bacillus*, *Brevibacterium*, *Corynebacterium*, *Enterococcus*, *Escherichia*, *Lactobacillus*, *Peptoniphilus*, *Prevotella*, *Staphylococcus*, and *Streptococcus* (**Figure 1B**). The most common species co-isolated with our clinical *P. mirabilis* strains were *Corynebacterium jeikeium*, *Enterococcus faecalis*, and *Staphylococcus epidermidis*.

**Figure 1.**
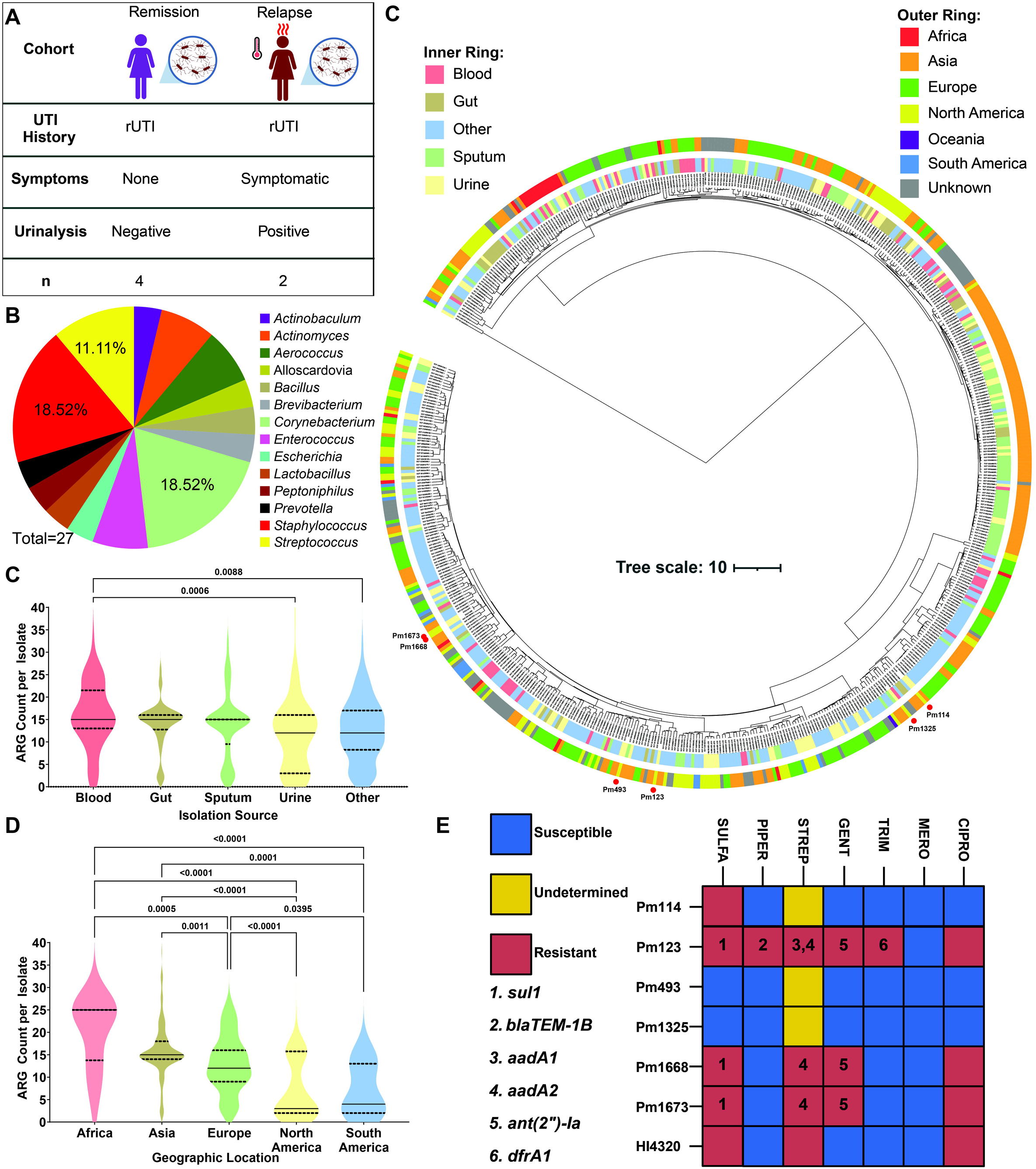
Isolation and genomic analysis of *P. mirabilis* isolates. A) Diagram showing cohort groups from which *P. mirabilis* strains were isolated. *Created in BioRender* (*2026*). https://BioRender.com B) Pie chart showing genera co-isolated with the six clinical *P. mirabilis* strains. C) Phylogenetic tree showing *P. mirabilis* strains from different isolation sources and geographic locations. Isolates from this study denoted by red circles. D) Violin plot showing ARG counts by isolation source, statistical analysis performed using Kruskal-Wallis test with Dunn’s multiple comparisons test. E) Violin plot showing ARG counts by geographic location. Oceania omitted as only one strain was from this region. Statistical analysis performed using Kruskal-Wallis test with Dunn’s multiple comparisons test. F) Resistance phenotypes for sulfamethoxazole (SULFA), piperacillin (PIPER), streptomycin (STREP), gentamicin (GENT), trimethoprim (TRIM), meropenem (MERO), and ciprofloxacin (CIPRO) with associated ARGs indicated by numbers. *Created in BioRender* (*2026*). https://BioRender.com

Following *P. mirabilis* strain isolation, Illumina and Oxford Nanopore sequencing was performed on the six clinical isolates and hybrid assemblies were generated using Unicycler v0.4.8 (41). All resulting genomes were found to be closed and circularized before being utilized for phylogenetic analysis (**Table 1**). To understand where our six rUTI-*associated P. mirabilis* strains fall in the greater *P. mirabilis* phylogeny, we downloaded publicly available *P. mirabilis* genomes from NCBI and 584 high-quality genomes were utilized in generating a phylogenetic tree of *P. mirabilis* isolates. The majority of genomes fell into one of two major clades, with a third small clade only containing two genomes. Our six clinical isolates (denoted by red dots) all fell into the same major clade but were relatively evenly distributed across the clade (**Figure 1C**). Interestingly, we did observe close clustering of Pm1673 and Pm1668, which were the only two of our clinical *P. mirabilis* isolates recovered from women with active, symptomatic rUTI. *P. mirabilis* genomes, including our isolates, did not cluster strongly by isolation source with urine, gut, blood and sputum isolates relatively well distributed across the phylogenetic tree, reflecting the generalist nature of *P. mirabilis* as a species (**Figure 1C**). We similarly did not observe strong clustering by geographic location, except for a single relatively large cluster of phylogenetically similar *P. mirabilis* isolates from Asia (**Figure 1C**).

**Table 1.**
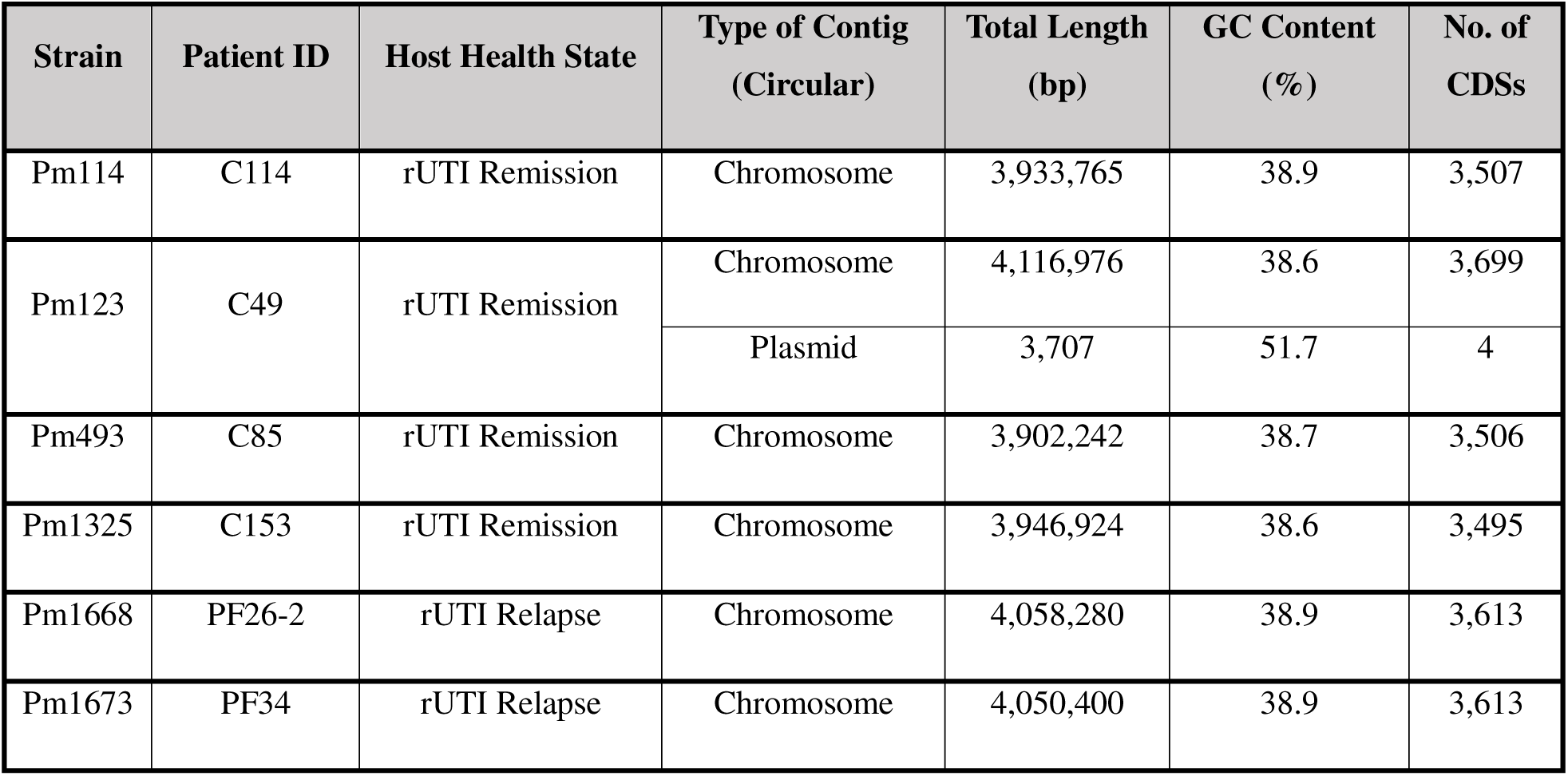
*Proteus mirabilis* genome assembly information.

To understand antibiotic resistance gene (ARG) carriage of our urinary *P. mirabilis* isolates and to determine any potential associations between ARG carriage and isolation source or geographic location, we performed ARG annotation of all genomes utilized in generation of the phylogenetic tree via ResFinder (42). The number of ARGs identified in an individual genome ranged from 2 to 38. The median ARG count among *P. mirabilis* blood isolates (15 ARGs) was significantly higher than the median ARG count for urine (12 ARGs) and other (12 ARGs) groups (**Figure 1D**). We also analyzed the number of ARGs per isolate by geographic region and found that isolates from Africa and Asia had a significantly higher median ARG count per isolate with a median of 25 and 15 ARGs per sample respectively as compared to Europe with a median of 12, North America with a median of 3, and South America with a median of 4 (**Figure 1E**). These data suggest a potentially lower incidence of MDR *P. mirabilis* in North and South America when compared to other continents. However, since resistance genotype may not always correspond with phenotype, we determined antibiotic minimum inhibitory concentration (MIC) for our clinical *P. mirabilis* strains as well as the HI4320 type strain. Interestingly, Pm123 had the most resistances, showing complete resistance to all tested antibiotics except for meropenem (**Figure 1F**). We also observed Pm493 and Pm1325 to have the weakest resistance profiles showing resistance to no tested antibiotics (**Figure 1F**). Importantly, in all cases where an ARG associated with a particular antibiotic resistance was identified, phenotypic resistance to the antibiotic was observed.

### Clinical *P. mirabilis* isolates exhibit differences in CS degradation and utilization

Prior research established chondroitin sulfate (CS) to be the most prevalent GAG in urine and that urinary *P. mirabilis* can break down CS and potentially use it as a carbon source (28, 29). We therefore sought to better understand how well conserved CS degradation was across the species and to examine differences in degradation efficiency. We first conducted a series of GAG degradation assays over a time course of 6, 8, 16, 24, and 48-hours for four of our clinical *P. mirabilis* strains, Pm123, Pm493, Pm1325, Pm1673, as well as the *P. mirabilis* type strain HI4320 in M9 minimal media supplemented with 0.75mg/mL of yeast extract (M9Ypm) to allow for growth of *P. mirabilis* (28). While CS degradation was observed for all tested *P. mirabilis* strains, differences in degradation kinetics were observed (**Figure 2A**). In terms of CS degradation activity in M9YPm, the strains fell into three groups of fast, moderate, and slow CS degraders. Fast CS degraders such as Pm493 and Pm1325 show the most rapid degradation and begin to break down CS within the first 6-hours of growth and degrade the majority of CS present within 8-hours. Moderate degraders HI4320 and Pm1673 degrade CS at an intermediate rate only beginning to break down CS after 8-hours with the majority being degraded after 16-hours of incubation (**Figure 2A**). Accordingly, both fast CS degrading strains, Pm493 and Pm1325, show more rapid growth stimulation by CS with growth stimulation over base M9Ypm observed by the initial 6-hour timepoint while the moderate strains only begin to show growth stimulation between 8 and 16-hours of incubation (**Figure 2B**). For the slow CS degrader, Pm123, substantial CS degradation was only observed after 24 hours and substantial growth stimulation was observed at 48-hours (**Figures 2A & 2B**).

**Figure 2.**
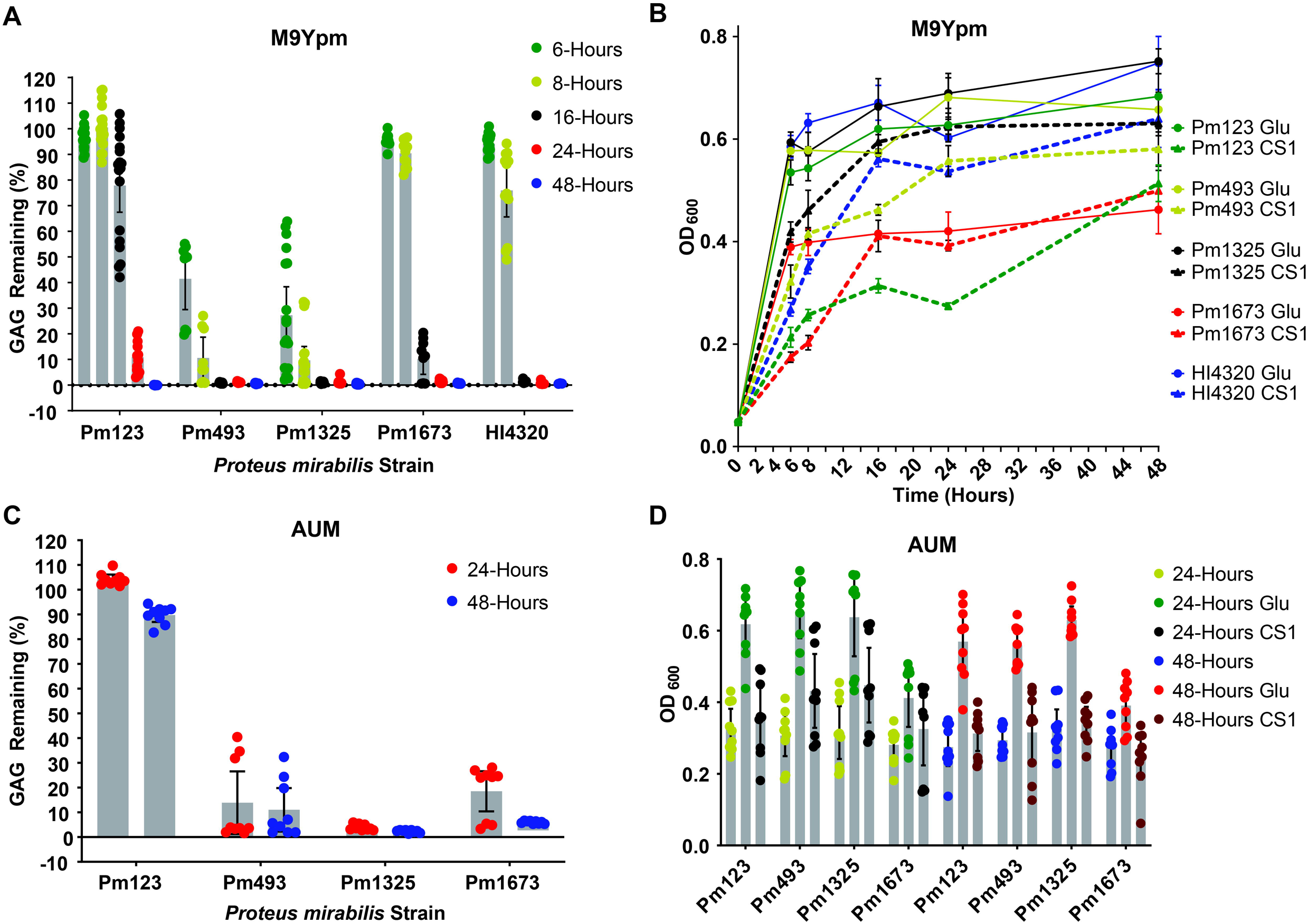
*P. mirabilis* strains exhibit variance in CS degradation kinetics in minimal and artificial urine media. A) 6 – 48-hour time-course of chondroitin sulfate (CS) degradation for clinical *P. mirabilis* isolates in M9Ypm. B) Growth of *P. mirabilis* isolates in basal M9Ypm versus with glucose or CS over 6 – 48-hours. C) Degradation of clinical *P. mirabilis* urinary isolates in AUM at 24 and 48-hours. D) Growth of *P. mirabilis* isolates in basal AUM versus with glucose or CS at 24 and 48-hours.

We next wanted to understand if our clinical *P. mirabilis* isolates would similarly degrade CS in a nutrient rich medium or if this activity was restricted to nutrient-limiting conditions. We therefore assayed CS degradation in Pm123, Pm493, Pm1325 and Pm1673 in Luria Bertani (LB) medium over a period of 24 to 48-hours. We observed robust CS degradation in LB after just 24 hours for Pm493, Pm1325 and Pm1673 (**Figure S1A**). The slow degrader, Pm123 was only able to degrade approximately 30% of the CS present in 24 hours but showed 100% degradation after 48 hours (**Figure S1A**). Endpoint growth assays showed that this degradation occurred in the absence of growth stimulation as the optical density for LB and LB + CS conditions were similar for all strains at 24 and 48 hours (**Figure S1B**). Interestingly, glucose seemed to reduce endopoint growth in LB for all strains at 24 hours and most noticeably for Pm1673 which exhibited greatly reduced growth in LB + glucose even after 48 hours (**Figure S1B**). We ultimately observed that while Pm123 degrades more slowly in LB, all other degradation phenotypes are comparable to that observed in M9Ypm with all other strains degrading fully within 24-hours and Pm123 fully degrading by 48-hours (Figures S2A & S2B).

Following the observation that *P. mirabilis* strains exhibit differences in CS degradation and utilization in minimal versus rich media, we wanted to assay CS degradation in a more physiologically relevant medium. As such, we conducted 24 and 48-hour CS GAG assays our four clinical *P. mirabilis* strains in artificial urine media (AUM) where it was observed that Pm493, Pm1325, and Pm1673 degrade the majority of CS present (∼60 – 100%) within 24-hours of incubation (**Figure 2C**). HI4320 was also found to degrade CS efficiently in AUM (**Figure S1A**). Interestingly, Pm123 was unable to degrade CS in AUM even after 48-hours of incubation (**Figure 2C**). All strains showed lesser growth stimulation by CS in AUM than observed in M9Ypm except HI4320 which showed similarly robust stimulation in both media types (**Figures 2D & S1B**). Pm123 showed absolutely no growth stimulation by CS in AUM even after 48-hours (**Figure 2D**).

**Urea represses chondroitin sulfate degradation by *P. mirabilis* Pm123**

To begin to understand the mechanism of repression Pm123 CS degradation in AUM, we sought to determine which component of AUM may be necessary for the observed phenotype. As urea is a major component of AUM that differentiates it from other media types and *P. mirabilis* encodes a well-characterized urease enzyme, we reasoned that urea would be a likely candidate (15–18). Upon removal of urea from AUM, Pm123 regained the ability to degrade urea similarly to what was observed in M9Ypm with nearly no GAG remaining after 24 hours. These data suggest that the presence of urea is necessary for repression of CS degradation by Pm123 (**Figure 3A**). In AUM with or without urea, no growth stimulation for Pm123 after 24 hours was observed (**Figure 3B**), but this is in line with our 24-hour observations in M9Ypm (**Figure 2B**).

**Figure 3.**
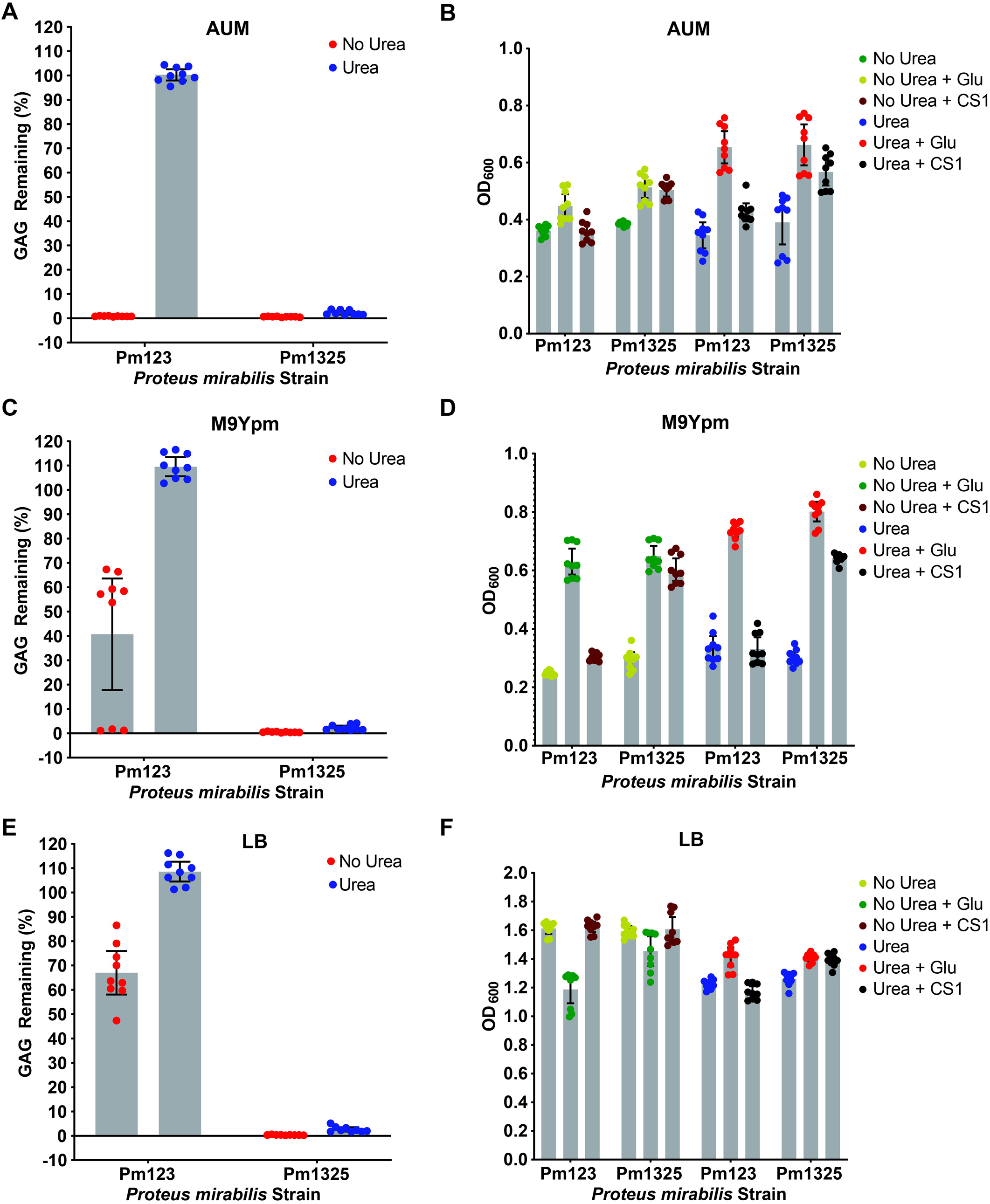
CS degradation in Pm123 is regulated by urea. A) Degradation of CS by Pm123 and Pm1325 in artificial urine media (AUM) with and without urea. B) Growth of Pm123 and Pm1325 in AUM with and without urea with either no carbon source, glucose, or CS. C) Degradation of CS by Pm123 and Pm1325 in M9Ypm with and without urea. D) Growth of Pm123 and Pm1325 in M9Ypm with and without urea with either no carbon source, glucose, or CS. E) Degradation of CS by Pm 123 and Pm1325 in LB with and without urea. F) Growth of Pm123 and Pm1325 in LB with and without urea with either no carbon source, glucose, or CS.

To determine if urea was sufficient to repress Pm123 CS degradation we supplemented M9Ypm with urea at the same concentration it is found in AUM (10mg/mL). After addition of urea, we observed a complete loss of CS degradation by Pm123 with the normal M9Ypm degrading ∼65% and the M9Ypm after 24 hours while the urea condition (M9YUpm) showed no degradation of CS for Pm123 but still complete CS degradation for the Pm1325 (**Figure 3C**). No substantial stimulation of Pm123 growth by was observed in either condition at the 24-hour time point, but CS stimulated Pm1325 growth in both urea and no urea conditions (**Figure 3D**).

Finally, as we see very little growth in M9Ypm or AUM with CS for Pm123 due to it being unable to utilize GAGs within the first 24-hours of growth, we tested if the urea repression phenotype could be replicated in a more nutrient rich media. We conducted a 24-hour GAG assay in LB compared to LB with 10mg/mL of urea (LBU). While we observed ∼60 – 80% GAG remaining after 24-hours in the no urea condition, like our previous observations in LB (**Figure S1**), there was no observed CS degradation for Pm123 in LBU (**Figure 3E**). This indicates that urea repression of Pm123 chondroitinase activity is not tied to lack of growth as all conditions showed growth of OD_600_ >1.0 and Pm123 still shows complete loss of CS degradation activity in the presence of urea (**Figure 3F**). As expected, CS degradation by Pm1325 in LB was unaffected by the addition of urea (**Figure 3E**). We also screened three additional *P. mirabilis* strains, Pm102-0, Pm104-0, and Pm106-15, isolated from CAUTI patients for potential urea inhibition of CS degradation and found that all three strains degrade CS regardless if urea is present but similarly to Pm1325 do not exhibit further growth stimulation by CS after 24-hours when grown in LB (**Figure S3A & S3B**).

### Urea regulation of CS degradation may be due to strain-level differences in *chABCI* function

To understand how urea may be regulating CS degradation in Pm123, we first sought to identify the enzymes contributing to CS degradation *P. mirabilis.* Through genome mining, we found that all *P. mirabilis* strains used in this study encoded homologs to the two well characterized *Proteus vulgaris* chondroitin lyases, *chABCI* endolyase and *chABCII* exolyase (38–40). To determine whether these genes are necessary for CS degradation in our clinical *P. mirabilis* strains, we generated markerless double knockouts of both genes in Pm123, Pm1668, and Pm1673 as we found these strains to be genetically tractable. Upon deletion of both *chABCI* and *chABCII* genes, we observed a complete loss of CS degradation by all three strains indicating that one or both of these conserved genes is required for CS degradation by our *P. mirabilis* strains (**Figure S4A**). Growth stimulation by CS was also lost in all double knockout strains (**Figure S4B**). With the putative chondroitinase genes identified, we next sought to determine if expression of either chondroitinase gene was repressed by urea in LB. We performed quantitative RT-PCR for the *chABCI* and *chABCII* genes of *P. mirabilis* grown in the presence or absence of urea and observed no statistically significant change in expression of either *chABCI* or *chABCII* after 4 hours of urea exposure for either Pm123 or Pm1325 (**Figure 4A**). We further tested if *chABCI* or *chABCII* expression was induced by CS addition for Pm123 and Pm1325 and observed no significant difference in expression of either gene in after four hours of CS exposure (**Figure 4B**). These results indicate that, at least within 4 hours of exposure, neither urea nor CS impact the expression of the *chABCI* or *chABCII* genes in Pm123 or Pm1325.

**Figure 4.**
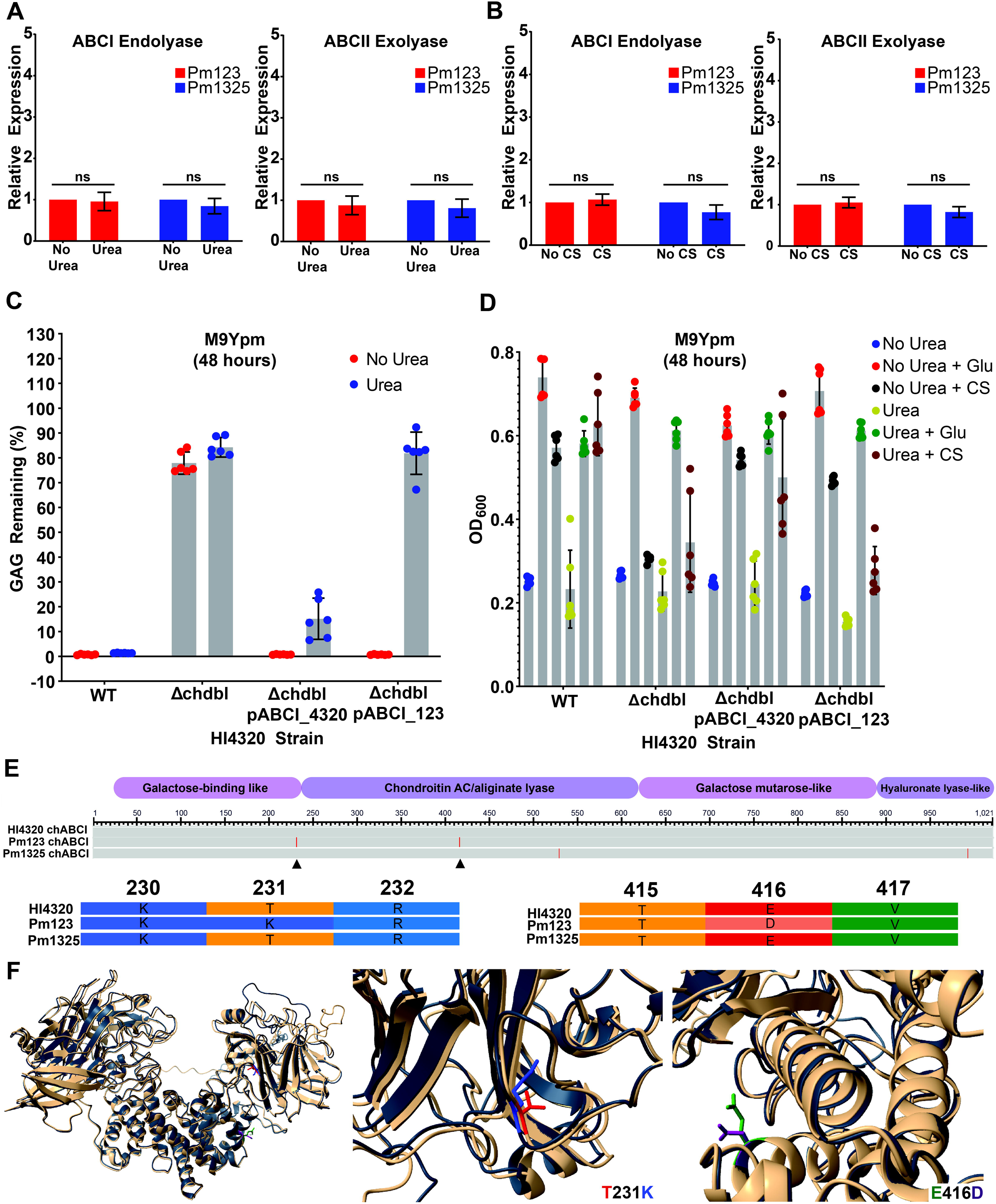
Urea regulation of CS degradation may be due to strain-level differences in *chABCI* function. A) Relative expression of *chABCI* and *chABCII* for Pm123 and Pm1325 grown with CS with or without urea for four hours. Statistical analysis performed using Mann-Whitney U test. B) Relative expression of *chABCI* and *chABCII* for Pm123 and Pm1325 grown with and without CS for four hours. Statistical analysis performed via Welch’s t-test as data passed normality test. C) Degradation of CS by HI4320 WT with empty pGEN vector, HI4320 *chabcI* and *chABCII* double knockout (Δchdbl) with empty pGEN, HI4320 Δchdbl with pGEN bearing the *chABCI* gene from HI4320, and HI4320 Δchdbl with pGEN bearing the *chABCI* gene from Pm123. D) Growth of HI4320 strains from panel C with and without urea with no carbon source, glucose, and CS. E) Protein sequence alignment of the chABCI enzyme from Pm123, Pm1325, and HI4320 with InterProScan predicted domains. F) 3D structural alignment of AlphaFold predicted chABCI enzyme structures for Pm123 (navy blue) and HI4320 (beige).

Next, to determine whether differences in the Pm123 enzyme structure or activity may be responsible for the urea sensitive phenotype, we compared the activity of *P. mirabilis chABCI and chABCII* double deletion (Δ*chdbl)* complemented either with the HI4320 *chABCI* endolyase or the Pm123 *chABCI* endolyase in the presence of urea. Without urea, HI4320 wild-type, HI4320Δ*chdbl* + *pchABCI_HI4320_,* HI4320Δ*chdbl* + *pchABCI_Pm123_* all degraded CS while HI4320Δ*chdbl* did not degrade CS indicating that the complemented chABC1 enzymes were functional (**Figure 4C**). However, upon the addition of urea, HI4320Δ*chdbl* cross-complemented with the Pm123 chABCI enzyme lost the ability to degrade CS while the strain complemented with the HI4320 chABCI enzyme was unaffected (**Figure 4C**). Accordingly, urea inhibited growth stimulation by CS only for HI4320Δ*chdbl* cross-complemented with the Pm123 chABCI enzyme (**Figure 4D**).

Since the cross-complementation experiments suggested potential differences in Pm123 and HI4320 chABCI enzyme function, we performed a sequence alignment of the chABCI endolyase protein sequences from HI4320, Pm123, and Pm1325 to identify any mutations unique to Pm123. We discovered two unique mutations in the Pm123 chABCI enzyme, the first being a threonine to lysine substitution at residue 231 and the other being a glutamic acid to aspartic acid substitution at residue 416 (**Figure 4E**) (43, 44). Domain prediction through InterProScan v5.77-108.0 revealed the T231K mutation to be in the N-terminal sugar binding domain and the D416E mutation to be in the chondroitin AC lyase domain. To pinpoint the location of these mutations in the 3D protein structure, we generated 3D structure predictions of each enzyme using AlphaFold 3 and performed a 3D structural alignment with ChimeraX v1.11.1 (**Figure 4F**) (45–48). The D416E mutation, although in a catalytic domain, was not localized to the catalytic cleft of the enzyme appearing to me more solvent exposed, while the T231K mutation was localized more internally in the sugar binding domain (**Figure 4F**).

### Urea inhibits chondroitinase activity through a urease dependent mechanism

We next sought to understand if urea was directly inhibiting the Pm123 chondroitinase activity or if, because *P. mirabilis* encode a well-characterized urease enzyme, the repression may be due to the breakdown products of urea. As such, we first performed a series of experiments to determine whether urease activity differs significantly between our *P. mirabilis* strains utilizing a phenol-red based urease assay and discovered that urea sensitive Pm123 and urea insensitive Pm1325 do not exhibit substantial differences in urease activity (**Figure 5A**). To test the hypothesis that urease activity contributes to urea repression of Pm123 chondroitinase activity, we first performed a dose-response experiment to determine the appropriate concentration of acetohydroxamic acid (AHA), a known urease inhibitor, needed to repress urease activity in both Pm123 and Pm1325. We determined 20mM AHA to be sufficient for inhibition of urease activity in both strains (**Figure 5B**). We next conducted a series of GAG assays in LB and LBU with or without AHA and observed that the addition of AHA restored the ability of Pm123 to degrade CS in the presence of urea even as soon as 24 hours (**Figure 5C**). Additionally, by 48-hours, we observed complete degradation of CS by Pm123 in LB, LB + AHA, and LBU + AHA while still showing near complete inhibition of Pm123 CS degradation in LBU with the 95% confidence interval showing between 80 and 110% GAG remaining after 48-hours (**Figure 5D**). These data suggest that inhibition of Pm123 chondroitinase activity by urea is tied to the action of the urease enzyme. The dependence of urea inhibition of Pm123 chondroitinase activity on urease activity suggests the inhibition is due to products of urea breakdown (i.e. ammonia, high pH) instead of the urea molecule directly.

**Figure 5.**
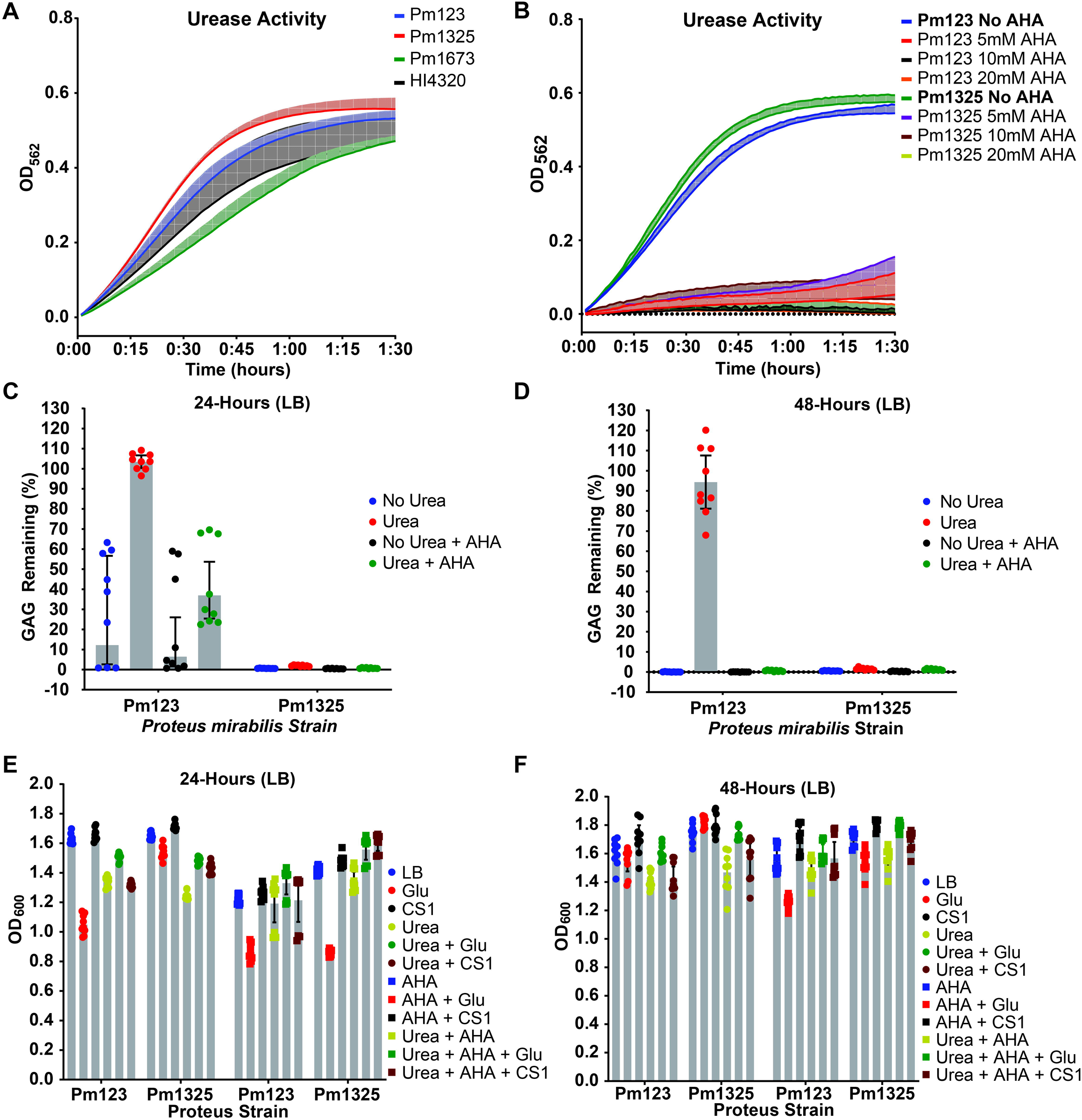
Urea inhibition of Pm123 CS degradation is dependent on *P. mirabilis* urease activity. A) Urease activity of the clinical *P. mirabilis* isolates compared to HI4320. B) Dose response of AHA inhibition of *P. mirabilis* urease activity for urea-sensitive Pm123 and urea-insensitive Pm1325. C) CS degradation for Pm123 and Pm1325 in LB with or without urea and with or without AHA at 24-hours. D) Pm123 and Pm1325 CS degradation in LB with or without urea and with or without AHA at 48 hours. E) 24-hour growth of *P. mirabilis* Pm123 and Pm1325 in LB with our without urea and with or without AHA in base media or with added glucose or CS. F) 48-hour growth of *P. mirabilis* with our without urea and with or without AHA in base media or with added glucose or CS.

### Virulence characteristics and the contribution of CS degradation to CAUTI repression differ between urea sensitive and insensitive *P. mirabilis* strains

We next wanted to understand if there were differences in virulence characteristics between our urea sensitive and urea insensitive *P. mirabilis* strains. We first assessed differences in swarming motility between our *P. mirabilis* isolates both in the presence and absence of urea (49). After measuring swarm colony diameter at 12 and 17-hours post-inoculation, we observed that Pm123 exhibits substantially less swarming motility than all other strains, with an average 12-hour swarm diameter of 25mm compared to 80-90 on average for the other strains (**Figure 6A**). Even after 17-hours the Pm123 swarm diameter only reached 35mm (**Figure 6B**). However, Pm123 swarming was not significantly impacted by urea either at the 12 or 17-hour time points. Comparatively, we observed that urea may impact HI4320 swarming with swarm colonies of ∼55mm after 12-hours with urea and ∼70mm with urea after 17-hours consistent with previous observations (**Figures 6A & 6B**) (50).

**Figure 6.**
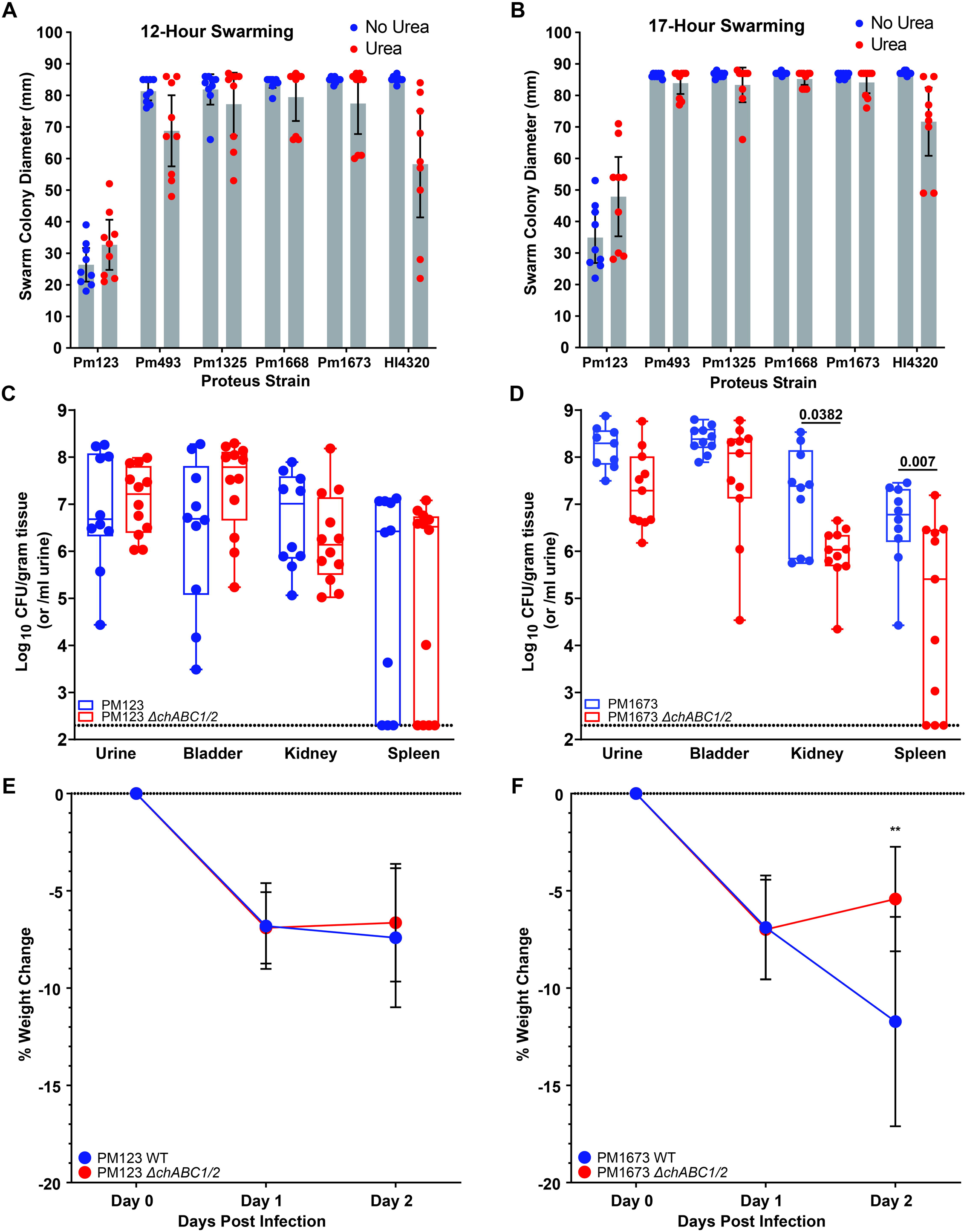
Differences in virulence traits and CS contribution to CAUTI between *P. mirabilis* strains. A) Swarm diameter for multiple *P. mirabilis* isolates after 12 hours. B) Swarm diameter for *P. mirabilis* isolates after 17 hours. C) CFU counts of Pm123 WT compared to Pm123 chABCI and II double knockout (ΔchABC1/2) in multiple tissues and urine after a 48-hour murine CAUTI model. Statistical analysis was performed by Kruskal-Wallis test with Dunn’s multiple comparisons correction as data were not normally distributed. D) CFU counts of Pm1673 WT compared to Pm1673 chABCI and II double knockout (ΔchABC1/2) in multiple tissues and urine after a 48-hour murine CAUTI model. Statistical analysis performed via parametric one-way ANOVA with Sidak’s multiple comparison correction as all samples passed a Kolmogorov-Smirnov normality test. E) Differences in mouse weight loss after CAUTI with either Pm123 WT or Pm123 ΔchABC1/2. Statistical analysis was performed by unpaired t-test with Welch’s correction. F) Differences in mouse weight loss after CAUTI with either Pm1673 WT or Pm1673 ΔchABC1/2. Statistical analysis performed using an unpaired t-test with Welch’s correction.

Finally, we aimed to determine if urea inhibition of Pm123 CS degradation would result in any difference in the contribution of chondroitinase activity to CAUTI disease outcomes. Using a well-established mouse model of CAUTI, we performed infections comparing wild-type urea sensitive Pm123 its isogenic *chABCI*/*II* double knockout and wild-type urea insensitive Pm1673 to its isogenic *chABCI*/*II* double knockout strain. Pm123 exhibited no statistically significant differences in disease outcome measured by CFU/g of bladder, kidney, and spleen or CFU/mL of urine between wild-type and the *chABCI*/*II* double knockout strain (**Figure 6C**). Conversely, for urea insensitive Pm1673, loss of chondroitinase activity was significantly associated with a reduction in kidney and spleen CFUs potentially suggesting a role for Pm1673 chondroitinase activity in progression of ascending UTI and invasion of other organs (**Figure 6D**). We observed a statistically significant decrease in weight of mice infected with wild-type Pm1673 as compared to its corresponding double *chABCI*/*II* knockout, but difference in weight was observed for mice infected with wild-type versus *chABCI*/*II* knockout Pm123 (Figures 6E & **6F**). These data indicate that while CS degradation contributes to virulence of urea-insensitive Pm1673 during CAUTI, it does not contribute to CAUTI virulence of urea-sensitive Pm123, highlighting potential *in vivo* implications of urea repression of Pm123 chondroitinase activity.

## Discussion

This study highlights the strain level diversity of CS degradation kinetics and regulation in the uropathogen *P. mirabilis.* Genomic analysis identified that all 6 clinical *P. mirabilis* strains encoded a putative chABCI endolyase and chABCII exolyase with high homology to the previously characterized *P. vulgaris* enzymes (37–40). All strains also encoded predicted phosphotransferase systems (PTS) downstream of the chondroitin lyases to potentially facilitate CS disaccharide import and utilization. However, despite the similarity among CS degradation and utilization gene clusters between strains, we observed major strain-level differences in the kinetics and regulation of CS degradation. Generally, *P. mirabilis* strains fell into three groups which we termed “fast”, “moderate”, and “slow” degraders with Pm123 exhibiting the slowest CS degradation of all strains in M9Ypm and LB. Pm1673 and HI4320 were observed as moderate strains showing slower degradation in M9Ypm than the two fast-degrading strains, Pm493 and Pm1325. The differences observed in M9Ypm indicate that CS degradation by different *P. mirabilis* strains varies greatly in nutrient-limited conditions. This may indicate that certain strains either take longer to “switch on” the cellular machinery required to break down CS or that differences exist in their enzymatic activity. The substantial delay exhibited by Pm123 in beginning to degrade CS and utilizing the breakdown products for growth possibly indicates that the switch in metabolism to utilizing CS breakdown products may be less efficient for certain strains or that they may preferentially breakdown other carbon sources. This hypothesis could also be supported by the slower CS degradation by Pm123 observed in LB when compared to M9YPm as it may be preferentially utilizing nutrients in LB before later switching over to CS as a carbon source. Ultimately, these differences highlight a need for a more detailed understanding of the regulatory pathways governing CS degradation and utilization in *P. mirabilis*.

This study provides insight into one potential mechanism by which chondroitinase activity may be regulated in the urinary tract. *P. mirabilis* strain Pm123 was unable to degrade CS in AUM due to the presence of urea. Removal of urea from AUM results in the restoration of CS degradation by Pm123 and conversely, addition of urea to the minimal media M9Ypm results in a complete inhibition of CS degradation. Urea repression of Pm123 chondroitinase activity was also independent of growth as we observed that urea also inhibited CS degradation in LB where high levels of growth were observed. We further determined that the urea repression of chondroitinase activity in Pm123 may be due to stain-level differences in the chABCI endolyase as complementation of a double *chABCI/II* knockout HI4320 strain with the *chABCI* gene from Pm123 conferred urea-sensitivity to HI4320 replicating the CS degradation phenotype observed in the Pm123 wild-type. We hypothesize that the urea-sensitivity of the Pm123 chABCI endolyase is a result of changes to enzymatic activity rather than differences in expression as urea did not impact gene *chABCI* gene expression and we observed two unique mutations in Pm123 chABCI sugar binding and catalytic domains not shared by urea-insensitive strains (47, 48). In Pm123, a mutation in the codon for residue 231 results in a threonine to lysine substitution, while another mutation in the codon for residue 416 results in a glutamic acid to aspartic acid substitution. Although either or both mutations may contribute to the urea sensitivity of the enzyme, we hypothesize that the lysine substitution in the sugar binding domain, may introduce destabilization charge interactions that influence substrate binding at high pH, which is known to occur as a result of the breakdown of urea by *P. mirabilis* (15–18). Lending credence to this hypothesis, we observed that urea repression of Pm123 CS degradation was dependent on the activity of the *P. mirabilis* urease enzyme as urease inhibition by AHA resulted in complete restoration of Pm123 CS degradation in the presence of urea.

Importantly, we also observed differences between urea sensitive and urea insensitive *P. mirabilis* strains in the contribution of chondroitinase activity to virulence in a mouse CAUTI model. While loss of chondroitinase activity in urea insensitive Pm1673 significantly decreased invasion of the kidneys and spleen as well as reduced weight loss in a mouse CAUTI model, this difference was not observed between Pm123 wild-type and chondroitinase knockout strains. One possible explanation for this observation is that the Pm123 chondroitin lyases are not active in the mouse urinary tract due to the presence of high concentrations of urea. Indeed, mouse urine has been shown to be highly concentrated with urea levels at ∼1,800 mmol/L (51, 52). Perhaps in the high urea environment of the mouse bladder, wild-type Pm123 is completely unable to degrade CS and therefore behaves similarly to the chondroitinase double knockout in terms of CS degradation.

Finally, phylogenetic analysis of our six clinical urinary isolates associated with rUTI history as well as publicly available genomes from NCBI highlights the generalist nature of *P. mirabilis* with little clustering observed by isolation source or geographic location (53). We did observe some clustering of isolate genomes from Asia, but many of these strains came from the same BioProject. In the analysis of ARG carriage and antibiotic susceptibility, our urea-sensitive, slow CS degrader, Pm123, was again found to be an outlier. Pm123 carried more ARGs and demonstrated phenotypic resistance to more antibiotics than the other five rUTI-associated isolates and HI4320. Pm123 was resistant to multiple classes of antibiotics including sulfonamides, β-lactams, aminoglycosides, and fluroquinolones and was only susceptible to meropenem out of the 7 drugs tested. The resistance to the β-lactam, piperacillin, was particularly interesting as Pm123 also exhibited a strong swarming defect and prior research has shown that swarming reduces resistance to β-lactam antibiotics (54). These results, although associative, may hint to a possible trade-off between swarming motility and antibiotic resistance.

This study demonstrates that the ability of *P. mirabilis* to degrade the prevalent urinary GAG, chondroitin sulfate, is widespread and a potentially clinically relevant virulence mechanism that promotes worse disease outcomes during CAUTI. This CS degradation activity is conserved across multiple strains; however, a wide range of phenotypes appear to exist amongst different isolates opening up new questions as to the regulatory pathways that control CS degradation by *P. mirabilis* both in vivo and in the environment. This study also provides insight into a possible mechanism by which CS degradation is regulated by urea and *P. mirabilis* urease activity via strain level differences in chABCI enzyme function. This study represents a major step towards greater understanding of *P. mirabilis* GAG degradation as well as its potential role in promoting invasion and colonization of the urinary tract.

## Materials and methods

### Strain isolation

Clinical *P. mirabilis* isolates used in this study were originally isolated from clean-catch mid-stream urine from postmenopausal women either with a history of rUTI but no active infection at the time of sample collection (rUTI remission) or history of rUTI with a symptomatic infection at the time of sample collection (rUTI relapse) (**Figure 1A**). Urine samples were collected from patients visiting the urology clinic at the University of Texas Southwestern Medical Center as part of institutional review board-approved studies STU 032016-006 and MR 17-120. Strains were originally isolated on either Chromagar Orientation or blood agar plates and isolated colonies were inoculated into either brain heart infusion (BHI) or M9 supplemented with 15g/L tryptone, 10g/L yeast extract, 20% glucose, 0.001g/L NAD, 0.0025g hemin, 0.25g/L L-cysteine-HCl, and 0.0025g/L vitamin K. Strains were then grown overnight before being glycerol stocked. Finally, species confirmation was performed via Sanger sequencing of the 16S rRNA gene amplified by the 8F and 1492R primer set (55). Three additional strains from catheterized individuals were obtained from an existing collection of bacteria isolates from the urine of nursing home residents with long-term catheters in Buffalo, New York between July 2019 and March 2020. Urine specimens were cultured on MacConkey agar and Columbia Nutrient Agar, and Gram-negative bacteria were identified to the species level using API-20E test strips (BioMérieux) (56). Species designation was further confirmed by whole genome sequencing (57).

### Bacterial strains and culture conditions

Clinical isolates and type strains of *P. mirabilis* were struck from glycerol stocks onto BHI agar plates to isolate single colonies. Individual colonies were then grown overnight in 3mL of BHI in glass culture tubes at 37 in ambient atmosphere and without shaking. For GAG assays, *P. mirabilis* was cultured in either M9 minimal media supplemented with 0.75mg/mL yeast extract (M9Ypm), artificial urine media (AUM), or LB (**Table S1**) (58). For acetohydroxamic acid (AHA) urease inhibitor assays, 50mM AHA was dissolved in milliQ H_2_O before being filter sterilized using a 0.2µm syringe filter.

### Bacterial genome assembly and phylogenetic analysis

Hybrid assemblies of *P. mirabilis* clinical isolates were prepared according to established protocols (59). Briefly, genomic DNA was first extracted using a Qiagen DNeasy blood and tissue kit. Oxford-Nanopore libraries were created using a ligation sequencing kit (SQK-LSK109) and barcode expansion kit 1-12 (EXP-NBD104) before being sequenced on a MinION device with R9 FLO-MIN106 flow cells. Illumina libraries were generated using a Nextera DNA Flex library prep kit and 2 x 150-bp paired-end reads were generated using a NextSeq 500 instrument. Short read quality assessment and trimming was then performed in CLC Genomics Workbench v12.0.3 and all reads with a length under 15bp and a Phred score under 20 were discarded. Finally, hybrid assemblies were generated using Unicycler v0.4.8 for all strains except Pm1325 which was generated using Unicycler v0.5.0, SPAdes v3.13.0, Racon v1.4.10, and Pilon v1.2.3 (41, 60–62). Genomes were then annotated using Prokka v1.14.6 for genome mining prior to online submission and the NCBI Prokaryotic Genome Annotation Pipeline v6.10 as part of genome submission to NCBI GenBank (63–66).

### Phylogenetic tree generation and antibiotic resistance gene prediction

Phylogenetic tree analysis was conducted using publicly available genomes from the NCBI genome database and categorized according to isolation source and location. In total, 584 *P. mirabilis* strains, including 64 blood isolates, 46 gut isolates, 101 sputum isolates, 105 urine isolates, and 268 isolates which either lacked relevant metadata or were collected from other sources passed inclusion criteria. Of the 584 isolates, 6 genomes were originally obtained as part of this study from patients in the Dallas, TX area. The remaining 578 isolates were downloaded from the NCBI genome assembly database on 20 January 2026 and included assembly levels from contig to complete, genomes were then filtered for a CheckM completeness value of ≧95 (67). Following genome collation and filtering, average nucleotide identity was calculated and a dendrogram was created with ANIclustermap v2.0.1 utilizing fastANI and seaborn all with default parameters (68–70). Finally, the resulting phylogenetic tree was visualized with iTOL v7 (71, 72). Following phylogenetic tree generation, antimicrobial resistance genes (ARGs) were predicted using ResFinder v4.0 from the Center for Genomic Epidemiology at the Technical University of Denmark with no provided species information, a minimum breadth-of coverage of 0.6, and identity threshold of 0.8 (42, 73).

### Antibiotic susceptibility testing

Strains were streaked on LB agar from frozen glycerol stocks and incubated overnight at 37°C. Individual colonies were used to inoculate Cation-Adjusted Mueller-Hinton Broth (CAMHB) which was then incubated overnight at 37°C and 200rpm. The following day, CAMHB subcultures were created using a 1:100 dilution and grown until exponential phase. Meanwhile, a two-fold dilution series was prepared using 10X stock solutions of the highest concentration to be tested of each antibiotic and then transferred to a 96-well plate, including a no antibiotic control well, to form triplicate rows. Subcultures were diluted to OD_600_ = 0.002 and 180 µL was added into the 96-well plate containing 20 µL of antibiotic dilution in each well. Plates were incubated at 37°C and 200rpm for 20 hours before OD_600_ measurement. MIC_90_ was determined by the sum of 10% of the growth measured in the no antibiotic control well with the background measurement of the plate. If the OD_600_ in the control well was greater than 1, it was treated as equaling to 1 to conservatively account for instrument inaccuracies at higher measurements. The lowest concentration well where the measured OD_600_ was below this threshold was determined to be the MIC.

### Semi-quantitative GAG degradation and utilization assay

The GAG degradation and utilization assay was performed as previously described (28). Briefly, overnight cultures were grown statically in BHI at 37 at ambient atmosphere, before being normalized to an OD_600_ of 0.05 in 1X PBS. Cells were then pelleted at 6010 x g for 8 minutes before being resuspended in assay liquid media. 300µL cultures were then aliquoted into 96-well plates in triplicate and plates were statically incubated in microaerophilic chamber (37, 5% O_2_, 10% CO_2_) for defined time periods. Following incubation, OD_600_ was recorded for all wells and then plates were centrifuged at 3214 x g for 10 minutes to pellet bacteria and 180µL of supernatant was removed into new plates. Serial dilutions were then performed in 1X PBS to the bottom of the established linear range for CS before 10µL of 10% bovine serum albumin (BSA) was added to all wells and OD_600_ was measured (pre-acetic acid control). Finally, 40µL of 2M acetic acid was added to all wells resulting in precipitation of the CS allowing for measurement of CS remaining using OD_600_. GAG remaining was then calculated according to the following formula using the OD_600_ value corresponding to the mid-point of the linear range for CS.

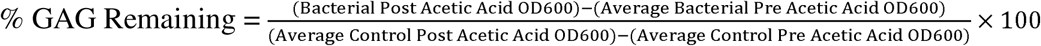

### Growth curves

Growth curves were conducted in LB with either no additional carbon source, glucose supplemented to 2.5mg/mL, or chondroitin sulfate supplemented to 2.5mg/mL to replicate conditions used in the semi-quantitative GAG degradation assay. Overnights were first grown in brain heart infusion (BHI) at 37 statically and then subcultured in LB to log phase (OD600 = 0.4 – 0.7). After reaching log phase, cultures were normalized to an OD_600_ of 0.05 in assay liquid media and aliquoted into 96-well plates. Growth curves were performed using a Cerillo Alto ALT-1600 inside of a microaerophilic chamber (37, 5% O_2_, 10% CO_2_) for 16-hours while shaking with readings taken every 5 minutes.

### Generation of *P. mirabilis* markerless deletion mutants

Markerless deletion mutants of clinical *P. mirabilis* urinary isolates were constructed via homologous recombination and *sacB* selection using the pDM4 vector (74, 75). First, homology regions ∼1kb upstream and downstream of the *chABCI* and *chABCII* genes were amplified via PCR (**Table S2**). Upstream amplicons were then digested with FastDigest restriction enzymes SalI and EcoRI, downstream amplicons were digested with FastDigest SacI and EcoRI, and the pDM4 vector was digested with SalI and SacI (ThermoFisher Scientific), for one hour and then ligated using Instant Sticky-end Ligase Master Mix (NEB) before being electroporated into *E. coli* S17 λpir. Cells were then allowed to recover at 37, shaking for one hour before being plated onto selection plates consisting of LB with 25µg/mL of chloramphenicol and grown overnight at 37 to isolate single colonies and verified by PCR. pDM4 chondroitin sulfate lyase knockout (CSKO) vectors were mated into *P. mirabilis* strains by first culturing *P. mirabilis* recipient strains in LB and *E. coli* donor strains in LB with 25µg/mL overnight at 37 shaking. Overnight cultures were normalized, washed in fresh LB and mixed at a 1:1 ratio in an Eppendorf tube. 100µL of mixed culture was then plated onto the center of LB agar plates containing 10mM MgCl_2_ to facilitate transfer of pDM4 CSKO vector and 2% agar to restrain *Proteus* swarming and incubated for 6 hours at 37. After mating, the bacterial lawn was scraped up with a pipette tip and resuspended in 1mL of sterile 1X PBS. Serial dilution was then performed down to 10^-3^ and dilutions were plated onto LB plates with 2% agar, 35µg/mL chloramphenicol and 20µg/mL tetracycline to select against the growth of the *E. coli* donor strain. Following positive selection, individual colonies were grown up overnight and then subjected to negative selection by plating on LB plates with 2% agar, 15% sucrose, and no NaCl. Following incubation overnight at 37, individual colonies were restruck onto negative selection media, incubated overnight at 37, and screened for *chABCI* and *chABCII* gene deletions by PCR.

The HI4320 chondroitinase double knockout strain was generated by the Armbruster lab by inserting a kanamycin resistance cassette into the gene of interest following the Sigma TargeTron group II intron protocol as previously described (76). Mutants were then screened by selection on kanamycin and PCR.

### Generation of HI4320 cross-complement strains

Complementation strains for the *chABCI* gene were generated in an HI4320 chondroitinase double knockout background (Δ*chdbl*) as follows. First, the *chABCI* gene, including 500bp upstream and downstream regions, was amplified by PCR from target HI4320 and Pm123 genomes using primers bearing the restriction sequences for the SmaI enzyme to enable the gene to be under native expression following insertion into the pGEN-MCS vector (77). The *chabcI* gene insert and purified pGEN vector were digested using FastDigest smaI (ThermoFisher Scientific) at 37 for 1 hour. Following digestion, the cut pGEN vector was treated with calf intestinal alkaline phosphatase (CIAP) (ThermoFisher) at 50 for 5 minutes. 10mM EDTA (pH = 8.0) was then added to the reaction mixture and CIAP was heat inactivated at 65 for 15 minutes. The insert was then ligated into the pGEN vector using T4 DNA ligase (NEB) overnight at 16, transformed into chemically competent DH5α *E. coli* cells, and transformants were selected on LB agar with 100µg/mL of ampicillin. Constructs were confirmed via restriction enzyme digestion with the SmaI enzyme. Successful *pGEN-chABCI* vector constructs were transformed into the HI4320 Δ*chdbl* strain via electroporation.

### RNA extraction and RT-qPCR

*P. mirabilis* strains Pm123 and Pm1325 were grown overnight in 3mL of BHI in glass culture tubes. Cultures were then normalized to an OD_600_ of 0.05 before being resuspended in 25mL of LB. Flask cultures were then grown in a hypoxia chamber (37, 5% O_2_, 10% CO_2_) for ∼16 hours before being normalized to OD_600_ = 0.25 in assay media (LB, LB + CS, or LBU + CS) and placed back into the hypoxia chamber and then harvested after 4-hours and treated with RNA Protect Bacterial Reagent (Qiagen). RNA was extracted using a Qiagen RNeasy Mini Plus kit and cDNA was prepared using qScript cDNA SuperMix. RT-qPCR was performed using PerfeCTa SYBR Green FastMix in a BioRad RT-qPCR machine and relative expression analysis was performed via the ΔΔCt method with expression normalized to the *recA* housekeeping gene. Statistical analysis was performed by conducting a Mann-Whitney U test on non-log transformed ΔCt values.

### Phenol-red urease assay

Urease activity was measured using a modified version of the whole-cell pH based urease assay protocol previously described (9, 78). Briefly, 3mL bacterial cultures were grown up overnight in glass culture tubes at 37 ambient with shaking. The next day, cultures were normalized to an OD_600_ of, pelleted and resuspended in 1mL of 1X PBS. 170µL of 1X PBS and 20µL of bacterial suspension was added into a 96-well plate followed by 10µL of phenol-red urea mixture (100µL of 0.1% wt/vol phenol red solution, 150mg of urea, 300µL of water, and 100µL of 1X PBS). A plate reader was used to collect OD_562_ readings every minute over a 2-hour kinetic interval with constant shaking.

### Swarming motility assay

Swarming motility assays were conducted as previously described (49). Briefly, 3mL bacterial cultures were grown overnight in LB at 37, shaking. The following day, cultures were normalized to an OD_600_ = 1 again in LB and 5µL of culture was dotted onto the center of agar plates containing 10g/L tryptone, 5g/L yeast extract, 10g/L sodium chloride, and 15g of agar (1.5%). Plates were incubated in a hypoxia chamber (37, 5% O_2_, 10% CO_2_) before swarm colony diameter was measured at 12 and 17-hours. Colony diameter was measured from the widest points at the edges of the swarm colony which bisected the center of original inoculum.

### Murine CAUTI model

Female CBA/J mice aged 6-8 weeks (Jackson Laboratory) were anesthetized with a weight appropriate dose of ketamine/xylazine (80-120mg/kg ketamine and 5-10 mg/kg xylazine) via IP injection and transurethrally inoculated with 50 µL of 2×10^6^ CFU/mL (1×10^5^ CFU/mouse) of PM123, a double chondroitinase knock-out mutant of Pm123, Pm1673 and a double chondroitinase knock-out mutant of Pm1673. A 4 mm segment of sterile silicone tubing (0.64 mm O.D., 0.30 mm I.D., Braintree Scientific Inc.) was advanced into the bladder during inoculation to be retained for the duration of the study as described previously (9, 79). After 48 hours, urine was collected, bladders, kidneys, and spleens were harvested and placed into 5 mL Eppendorf tubes containing 1 mL 1x PBS and 500 µL of 3.2mm stainless steel beads. Tissues were homogenized using a Bullet Blender 5 Gold (Next Advance, Speed 8, 4 minutes). Bladders were treated to two cycles to ensure full homogenization. Tissue homogenates were serially diluted and plated onto appropriate agar using an EddyJet 2 spiral plater (Neutec Group) for determination of CFUs using a ProtoCOL 3 automated colony counter (Synbiosis).

## Acknowledgements

We thank the University of Texas at Dallas Genome Center for their services and expertise. This work was supported by the Welch Foundation, award number AT-2030-20200401 to N.J.D., by the National Institutes of Health, award number R01DK136875 to C.E.A. and N.J.D, and by the Felecia and John Cain Distinguished Chair in Women’s Health, held by P.E.Z.

