## Supplemental Material for "Strain level variation in *Proteus mirabilis* chondroitin sulfate degradation kinetics and regulation by urea"

**Table S1. Formulation of media used for *P. mirabilis* GAG assays**

| **Media Type** | **Component** | **Quantity** | **Unit (per Liter)** |
| --- | --- | --- | --- |
| M9Ypm | 5X M9 Salt Solution | 200 | mL |
|  | 1M Magnesium Sulfate Heptahydrate | 2 | mL |
|  | 1M Calcium Chloride Dihydrate | 0.1 | mL |
|  | Yeast Extract | 0.75 | g |
| AUM | Peptone L37 | 1 | g |
|  | Yeast Extract | 0.005 | g |
|  | Sodium Bicarbonate | 2.1 | g |
|  | Sodium Chloride | 5.2 | g |
|  | Disodium Sulfate Decahydrate | 3.2 | g |
|  | Potassium Dihydrogen Phosphate | 0.95 | g |
|  | Dipotassium Hydrogen Phosphate | 1.2 | g |
|  | Ammonium Chloride | 1.3 | g |
|  | L-Lactic Acid | 0.1 | g |
|  | Urea | 10 | g |
|  | Citric Acid | 0.4 | g |
|  | Creatinine | 0.8 | g |
|  | Iron (II) Sulfate Heptahydrate | 0.0012 | g |
|  | Uric Acid | 0.07 | g |
|  | Calcium Chloride Dihydrate | 0.37 | g |
|  | Magnesium Sulfate Heptahydrate | 0.49 | g |
| LB | Tryptone | 10 | g |
|  | Yeast Extract | 5 | g |
|  | Sodium Chloride | 10 | g |

**Table S2. PCR primers list**

| **Primer name** | **Primer use** | **Primer sequence (5’ – 3’)** |
| --- | --- | --- |
| CSKO_Upstream_F | 1kb upstream region with SalI site for generation of ABCI/ABCII knockout in Pm123, Pm1668, and Pm1673 | GATCGTCGACTGCACTATCAAATGATCTGGCG |
| CSKO_Upstream_R | 1kb upstream region with EcoRI site for generation of ABCI/ABCII knockout in Pm123, Pm1668, and Pm1673 | GATCGAATTCTATATTTTCTCCTTAGAAACGCTGG |
| CSKO_Downstream_F | 1kb downstream region with EcoRI site for generation of ABCI/ABCII knockout in Pm123, Pm1668, and Pm1673 | GATCGAATTCAAGTCGTAACTTTTAAATTAAAGAGTC |
| CSKO_Downstream_R | 1kb downstream region with SacI site for generation of ABCI/ABCII knockout in Pm123, Pm1668, and Pm1673 | GATCGAGCTCGAGCATAATATGGATTATGGCGC |
| chABCI_qPCR_F | qPCR for chABCI endolyase | CTGGCGTGCTATTGGTATCT |
| chABCI_qPCR_R | qPRC for chABCI endolyase | CTGTCAACATTCGCACCTAAAG |
| chABCII_qPCR_F | qPCR for chABCII exolyase | GCTAAAGGGCAAACGGTAGA |
| chABCII_qPCR_R | qPCR for chABCII exolyase | GGGCTTTACCGTGAGAGATAAC |
| recA_qPCR_F | qPCR for recA | TCCGTGGCAGCATTAACA |
| recA_qPCR_R | qPCR for recA | TACGCATAGCTTGGCTCATC |
| ABCI_pGEN_F | Amplification of chABCI endolyase in Pm1325 with SmaI restriction site for cloning into pGEN | GATCCCCGGGCAAAGCAAAAGGTGACTACTCTG |
| ABCI_pGEN_R | Amplification of chABCI endolyase in Pm1325 with SmaI restriction site for cloning into pGEN | GATCCCCGGGCAGGAATAGGCCAACGATTATC |

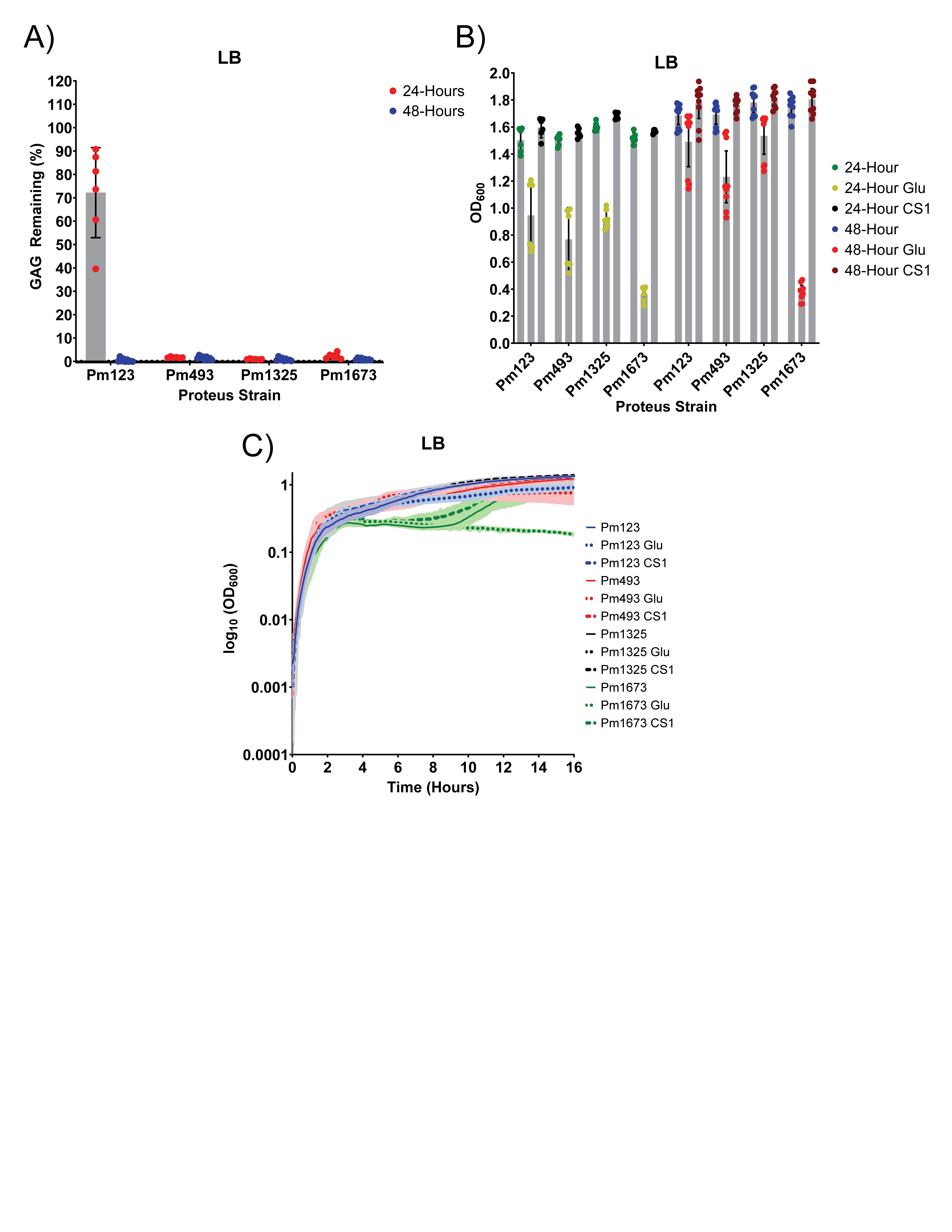

**Supplemental Figure 1. Degradation of CS and growth in LB**

A) Degradation of CS in LB by *P. mirabilis*. B) Growth of *P. mirabilis* in LB with no added carbon source, glucose, and CS. C) Growth curves of *P. mirabilis* strains in LB with no added carbon source, glucose, and CS.

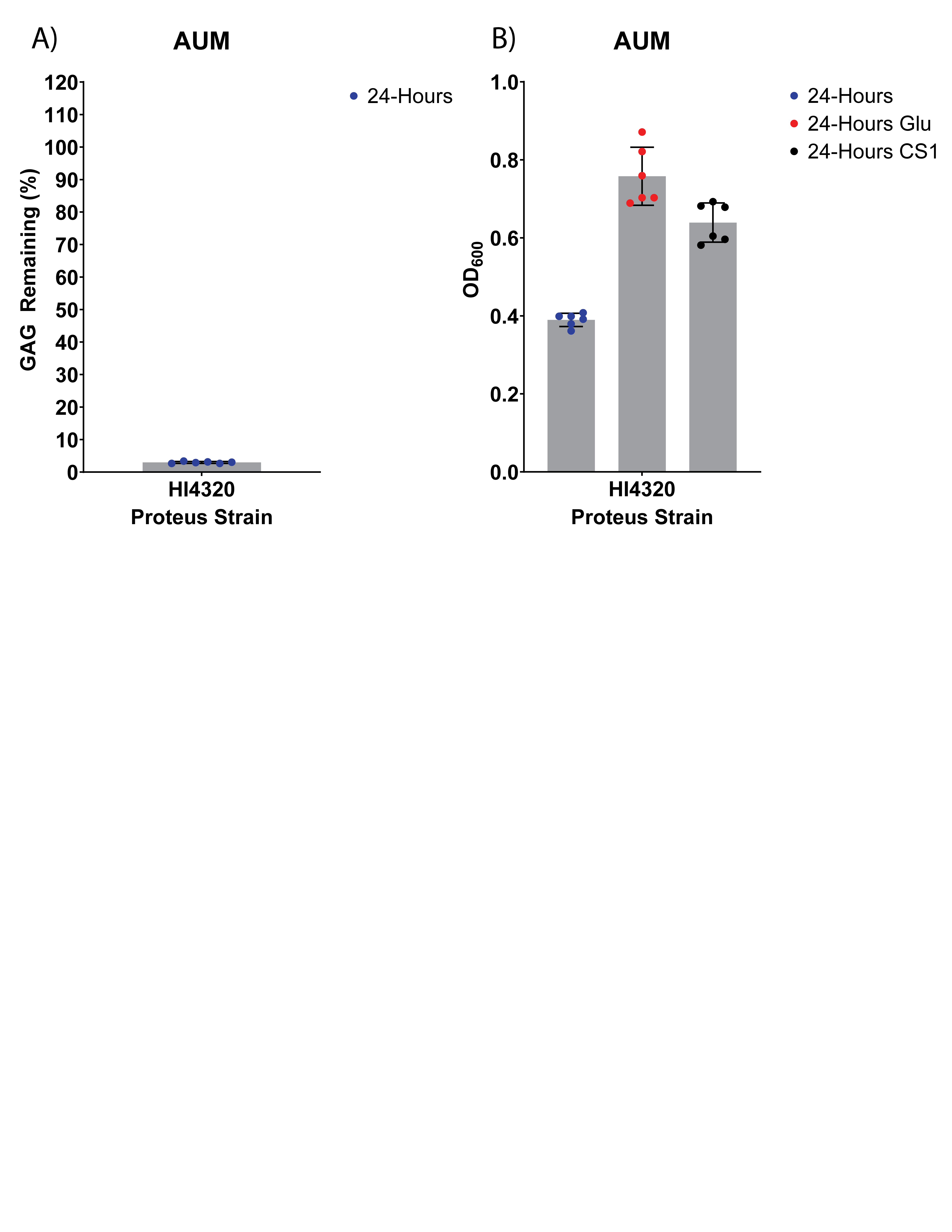
**Supplemental Figure 2. Degradation and utilization of CS by HI4320 in AUM**

A) Degradation of CS by HI4320 in AUM after 24 hours. B) Growth of HI4320 in AUM with no carbon source, glucose, or CS

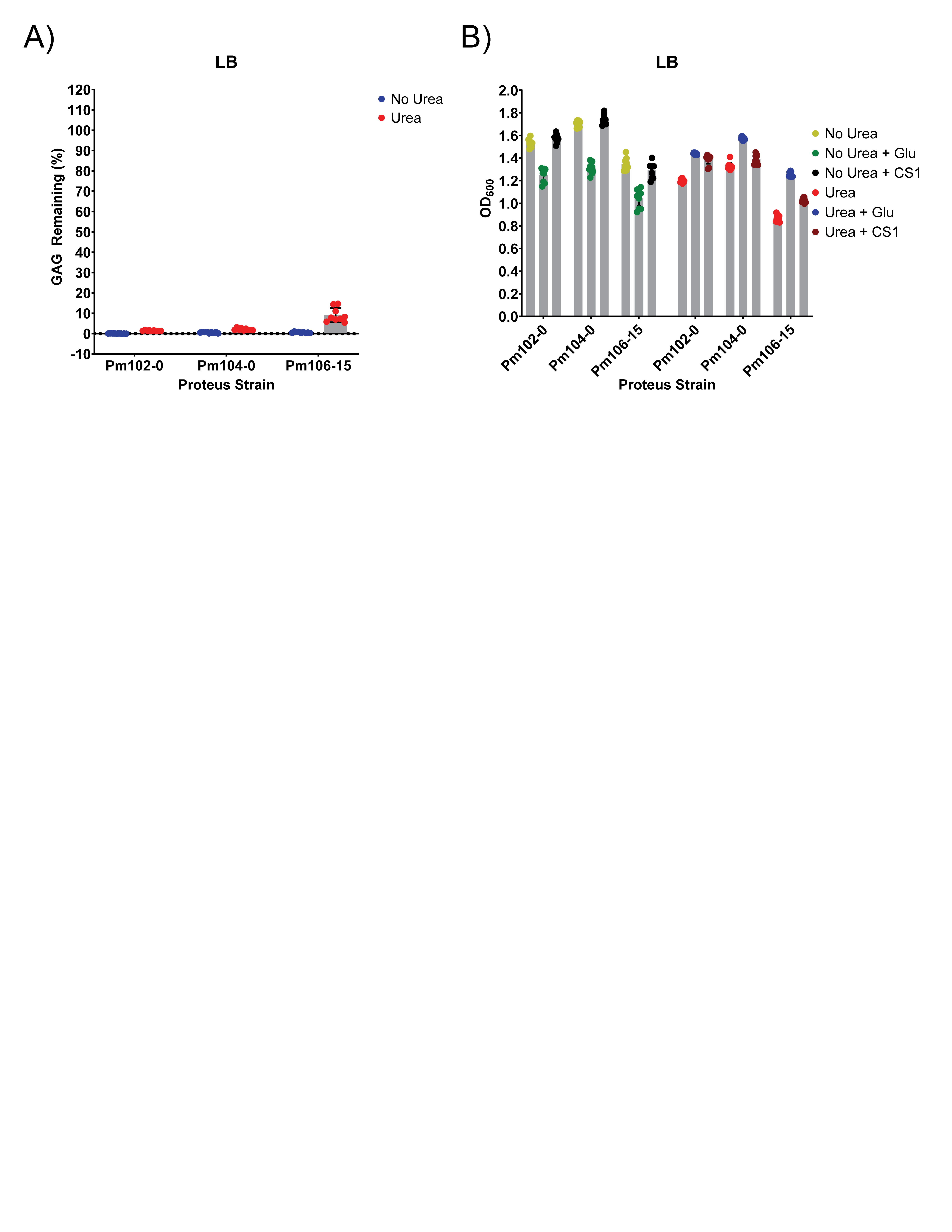

**Supplemental Figure 4 Degradation profiles of additional clinical *P. mirabilis* urinary isolates**

A) Degradation of CS by *P. mirabilis* clinical urinary isolates Pm102-0, Pm104-0, and Pm106-15 in LB and LBU. B) Growth of Pm102-0, Pm104-0, and Pm106-15 in LB and LBU with no additional carbon source, glucose, or CS.

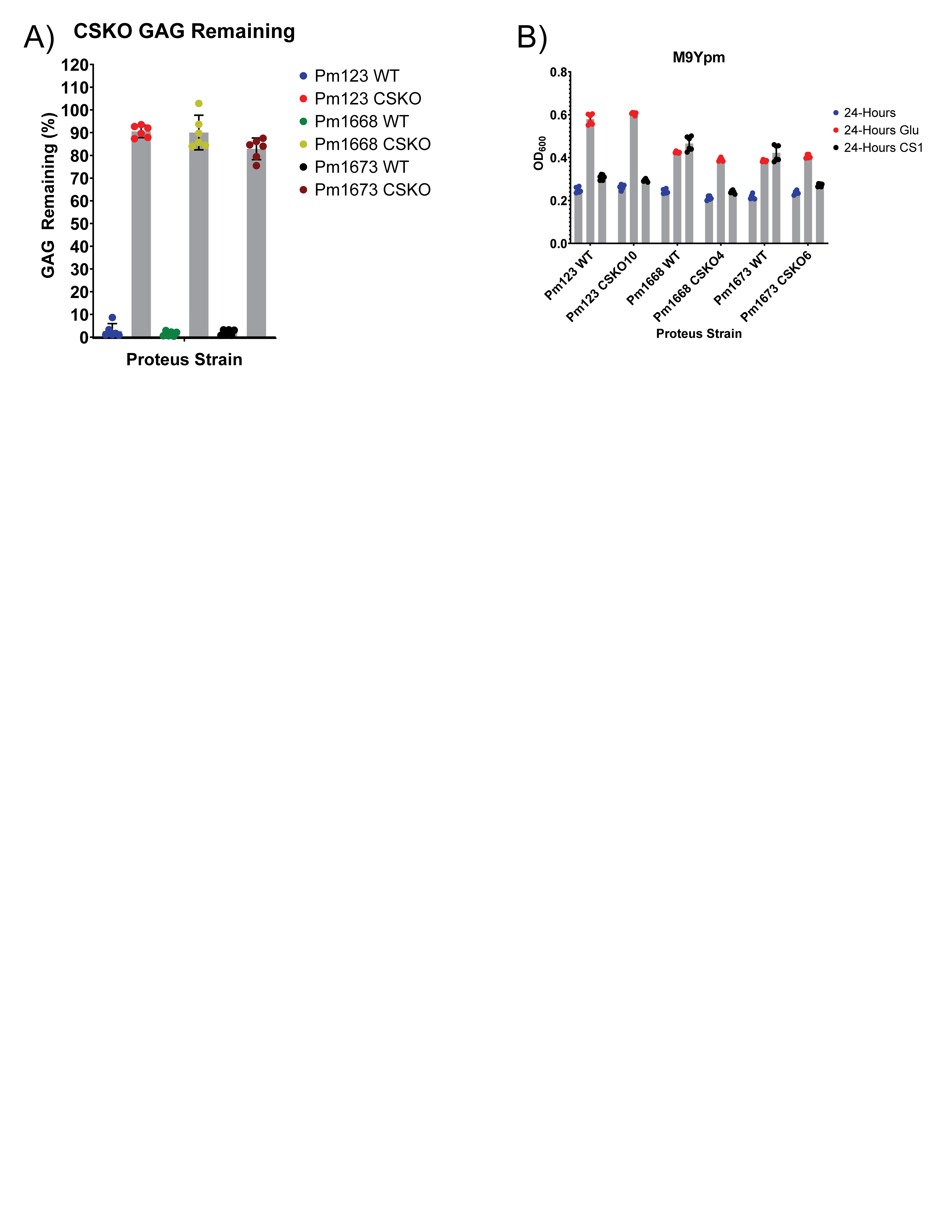

**Supplemental Figure 3 Loss of putative chondroitinase genes removes CS degradation** **ability**

A) CS degradation in M9Ypm of *P. mirabilis* strains Pm123, Pm1668, and Pm1673 WT compared to chondroitinase double knockout (CSKO). B) Growth of *P. mirabilis* strains Pm123, Pm1668, and Pm1673 WT vs CSKO strains in M9Ypm with no carbon source, glucose, or CS.
